# Serial, as opposed to parallel, insular-prefrontal cortex processing determines the tendency to make risky decisions

**DOI:** 10.64898/2026.06.25.734465

**Authors:** Dhaval D. Joshi, Kshitij Jhadav, Lanfei Sun, Tristan Hynes, David Belin

## Abstract

Adaptive decision-making under ambiguity requires constant integration of reward- and loss-related information to guide behaviour. In humans and rodents, not all individuals maximise gains in decision-making tasks, such as the Iowa Gambling task or its rodent version, the Rat Gambling task (rGT). While the prefrontal and insular cortices have each been shown independently to support optimal probabilistic decision-making, how they interact functionally to shape individual differences in performance remains unclear. Here, we investigated the consequences of bilateral baclofen/muscimol-mediated inactivation of the prelimbic cortex (PLC) or the anterior insular cortex (AIC) vs. their functional disconnection on the performance of Sprague Dawley rats identified as safe (SDMs) or risky decision makers (RDMs) in the RGT. AIC inhibition decreased advantageous choice in SDMs, whereas it increased win-stay responding in RDMs. In contrast, PLC inhibition primarily affected lose-shift behaviour, reducing sensitivity to losses in SDMs while enhancing adaptive switching in RDMs. Functionally disconnecting the PLC from the AIC, which had no effect on the performance of SDMs, improved decision-making in RDMs by increasing loss-guided behavioural adaptation. Together, these findings identify parallel versus serial AIC-PLC processing as a potential neural mechanism underlying the tendency some individuals have to make suboptimal decisions.

## Introduction

Success in daily life depends on the ability to make optimal decisions in ambiguous situations by learning the outcomes associated with available alternatives. This process is partially driven by reinforcement learning, whereby individuals repeat choices that yield rewards and avoid those that result in losses. Importantly, real-world decision-making is often probabilistic: advantageous choices do not always yield positive outcomes, while disadvantageous choices can occasionally yield large rewards. For example, one route home may typically be faster because traffic is usually lighter, yet unpredictable traffic patterns can occasionally make the alternative route more efficient. In humans [1–4] and non-human mammals, such as rodents, individuals vary substantially in their ability to make optimal decisions in such ambiguous environments [5–8], and severe impairments in this capacity are associated with substance use disorder and gambling disorder [9–12].

Decision-making under uncertainty can be assessed clinically and experimentally using the Iowa Gambling Task [IGT, 13], which requires participants to repeatedly choose between probabilistic options that offer high immediate rewards coupled with larger delayed cumulative losses, and options that yield smaller immediate gains but superior long-term outcomes [12]. Combined with neuropsychological characterisation of patients with lesions encompassing distinct brain territories or correlational brain imaging investigations, the IGT has been pivotal over the past 30 years in identifying the psychological and neurobiological substrates of decision-making under uncertainty [for a review, see 14]. Of these, the ventromedial prefrontal cortex [vmPFC, 15,16,17] and the insula [18–20] have been shown to play complementary roles in the processes, mostly implicit, that underlie decision-making performance, as outlined in the somatic marker hypothesis [21].

The insula computes risk predictions and guides choice in risk-sensitive individuals [22,23], likely by integrating interoceptive signals [24,25] that the prefrontal cortex subsequently uses for higher-order computations [24]. The vmPFC thus integrates information from the insula, along with multiple parameters of future outcomes, into a combined value [26], ultimately leading to the avoidance of suboptimal choices [13]. This function is considered to depend on the vmPFC’s ability to engage representations of possible outcomes and their associated autonomic-system-dependent emotional states [27,28].

A rodent analogue of the IGT – the rat gambling task (rGT) – has provided support for the causal roles of the PFC and the insula in probabilistic decision making [5,29]. In rats, pre-training bilateral excitotoxic lesions of the prelimbic cortex (PLC), a putative functional analogue of the vmPFC in humans [30], profoundly reduce the ability to choose advantageous options in the RGT [29], a deficit similar to that observed in patients with vmPFC lesions who persist in selecting immediately rewarding yet disadvantageous options despite accumulating losses [13,15,16]. In contrast, bilateral excitotoxic lesions of the anterior insular cortex (AIC), which shares structural, connectivity and functional features with the anterior portion of the insula in humans [31–33], result in a bidirectional effect on performance in the RGT [5]. Thus, lesioning the AIC impairs the performance of good decision-making individuals (safe decision makers, SDM), while symmetrically improving that of those who tended spontaneously to stick with riskier choices (risky decision makers, RDM). These observations revealed an important role for the AIC in individual differences in decision-making, the neural basis of which remains poorly elucidated.

Since the AIC forms an interconnected circuit with the PFC and both guide decision-making under ambiguity, we hypothesised that inter-individual differences in the functional organisation of this circuit underlie tendencies to engage in safe vs risky decision-making. We therefore compared the effects of bilateral reversible inactivation of either the PLC or the AIC to that of their functional disconnection on the performance of male rats identified as safe or risky decision-makers (SDMs and RDMs, respectively) in the RGT [5,34].

## Materials and methods

### Subjects

Seventy-two male Sprague Dawley rats (Charles River, UK), weighing 300-350 g at the start of experiments, were single-housed under a reversed 12h light/dark chain (lights off at 7:00 AM) and food restricted to gradually reach 85% of their free-feeding body weight before starting the behavioural training. Water was always available ad libitum. Experiments were performed 6-7 days/week between 8 AM and 5 PM. All experimental protocols were conducted under the project license 70/8072 held by David Belin in accordance with the regulatory requirements of the UK Animals (Scientific Procedures) Act 1986, amendment regulations 2012, following ethical review by the University of Cambridge Animal Welfare and Ethical Review Body AWERB.

### Rat Gambling Task

All rats were trained in the rGT as described previously [5,34].

Chiefly, testing took place in eight standard 5-hole operant chambers (Med. Associates, St Albans, VT, USA), each located within a sound-attenuating chamber equipped with exhaust fans that ensured air renewal and masked background noise. Operant chambers were illuminated by a white 3-W house light during experimental sessions. A pellet dispenser delivered 45-mg dustless precision pellets (Bio-Serv Inc., NJ, USA) to a food magazine on the right wall. On the opposite wall, the 5-hole stimulus array was positioned 2 cm above a bar floor, and each aperture contained a stimulus light. Nosepokes into the food magazine or the holes were recorded with infrared photobeams. The boxes were controlled by software written in MED-PC (Med. Associates) on a computer running under Windows 7.

Animals were exposed to sucrose pellets in their home cages before being habituated to the testing boxes. During initial magazine training, 60 pellets were delivered to the magazine on a 30-s variable interval schedule. Then, rats were trained to nose-poke into one of the four laterally illuminated holes to receive a food pellet reward. Responses in the middle, inoperative hole were recorded but had no programmed consequence. Sessions continued until rats obtained 100 pellets or 30 min elapsed.

After two free-choice training sessions, rats were given four forced-choice 30-minute sessions during which one of the four holes was active for 7 minutes and 30 s on a pseudorandom schedule. Forced-choice sessions were implemented to help animals avoid developing a side or hole bias. Subsequently, rats underwent two 30-minute free-choice sessions during which each nose poke in any of the four active holes provided two pellets during the first half of the session and one pellet during the second half or vice versa.

During the rGT proper, two of the four active holes, defined as the disadvantageous holes, resulted in the guaranteed delivery of two pellets but a potential long timeout of either 222 or 444 s, with probabilities of 0.5 and 0.25, respectively. The other two active holes, deemed advantageous, were instead associated with the delivery of only one pellet, potentially followed by a short timeout of either 6 or 12 s, with respective probabilities of 0.5 and 0.25. The probability of receiving a timeout punishment for each hole was fixed for the duration of the session. Options were distributed randomly across the 4 active holes of the chamber. The test session lasted until rats obtained 250 pellets or 90 min had elapsed.

As the utility within each pair of options was similar, choice of both advantageous options was pooled, as were choices from either disadvantageous option, to generate a decision-making score for each animal.

Rats in the top tercile of the overall percentage of safe decisions were designated as the safe decision makers (SDMs; n = 24). Those in the bottom tercile were designated risky decision makers (RDMs; n = 24). Rats in the intermediate tercile (n = 24) were used in a different experiment. Following behavioural phenotyping, rats were randomly assigned to surgical groups for (n = 8 / phenotype / manipulation).

### Intracranial Surgery

SDMs and RDMs underwent stereotaxic surgery under general anaesthesia (isoflurane, 5% induction / 2% maintenance) to bilaterally implant guide cannulae (22-gauge, Plastics One, Roanoke, VA, USA) that terminated 2 mm above the PLC (coordinates from bregma: anterior/posterior -AP- +3.3) mm, mediolateral (ML) ± 0.8 mm, dorsal/ventral (DV) -3.8 mm from skull) (experiment 1, **Figure 1**), the AIC (AP +1.44) mm, ML ±5.3 mm, DV (-4.9 mm from skull) (experiment 2, **Figure 1**, or both (**Figure 3**). Cannulae were held in place using dental acrylic cement anchored to four stainless steel screws tapped into the frontal and parietal bones of the skull. Obturators (Plastics One, Roanoke, VA, USA) were placed in the cannulae to maintain patency. All rats were given five days to recover from surgery, and for the first three days after surgery, they were treated daily with 1mg/kg anti-inflammatory drug Metacam (Boehringer Ingelheim, Ingelheim am Rhein, Germany) orally administered in drinking water.

**Figure 1.**
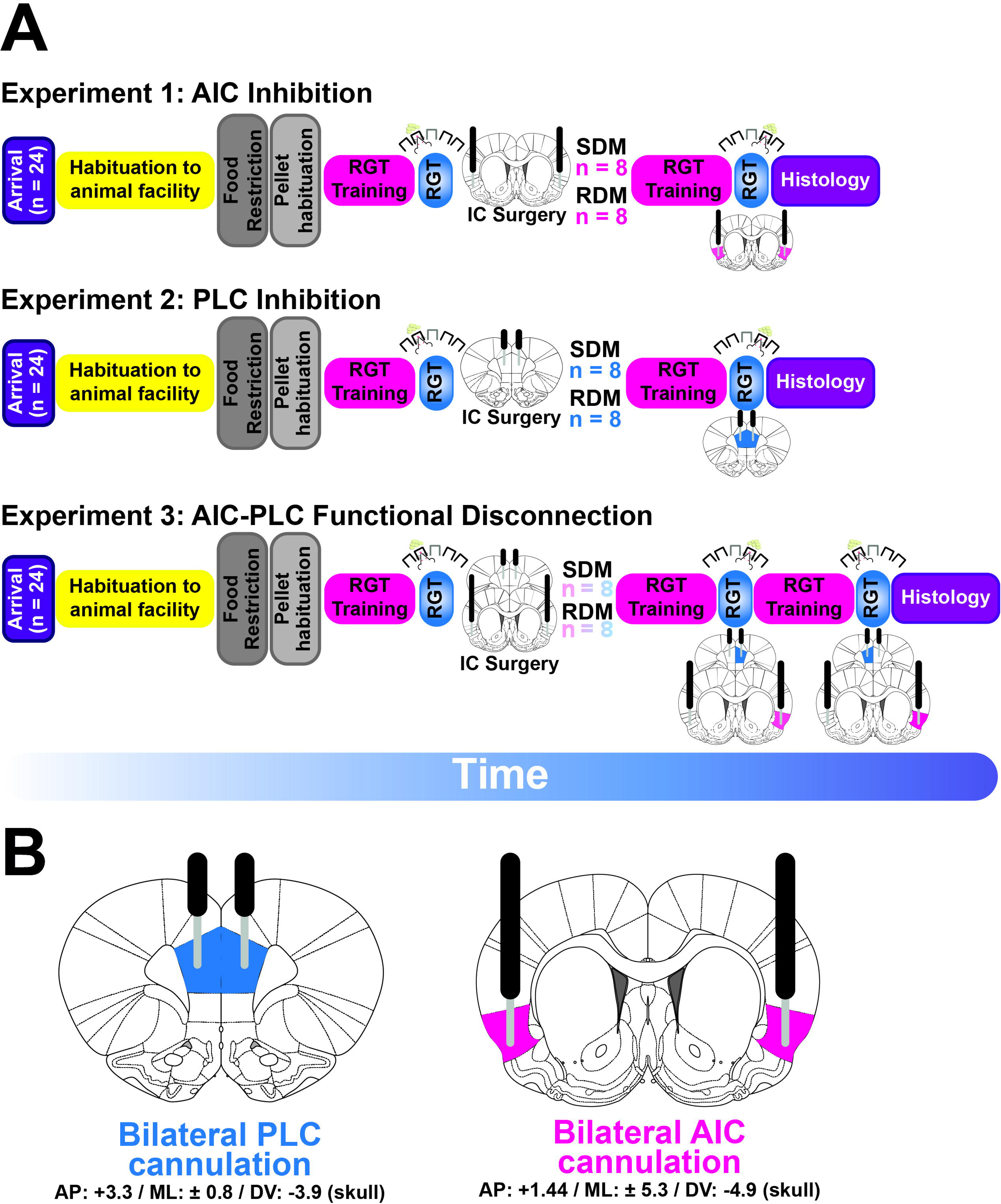
Schematic of experimental design. (A) Timelines of the three independent studies. Experiments 1, 2, and 3 aimed to address the functional roles of the anterior insular cortex (AIC), the prelimbic cortex (PLC), and AIC-PIC interactions in risky decision-making under ambiguity, respectively. In each experiment, rats were trained on the RGT. Following the characterisation of Risky- (RDMs) and Safe-decision makers (SDMs), each phenotype group underwent intracranial bilateral cannulation surgeries of the AIC (Exp. 1), PLC (Exp. 2) or both (Exp. 2). In Experiments 1 and 2, following recovery from surgeries, the risky decision-making was re-assessed following inactivation via baclofen/muscimol infusion into the AIC or PLC, respectively. In Experiment 3, animals were tested twice following cannulation. Once following ipsilateral inactivation of the AIC and PLC, and once following contralateral inactivation (the order of inactivation manipulation was counterbalanced between animals). After the final RGT test, all animals were transcardially perfused and their brains were processed for cresyl staining to ascertain cannula placement. (**B**) Displays the positioning of the guide (black) and injector (grey) cannulae into the two cortical territories along with the stereotaxic coordinates.

**Figure 2.**
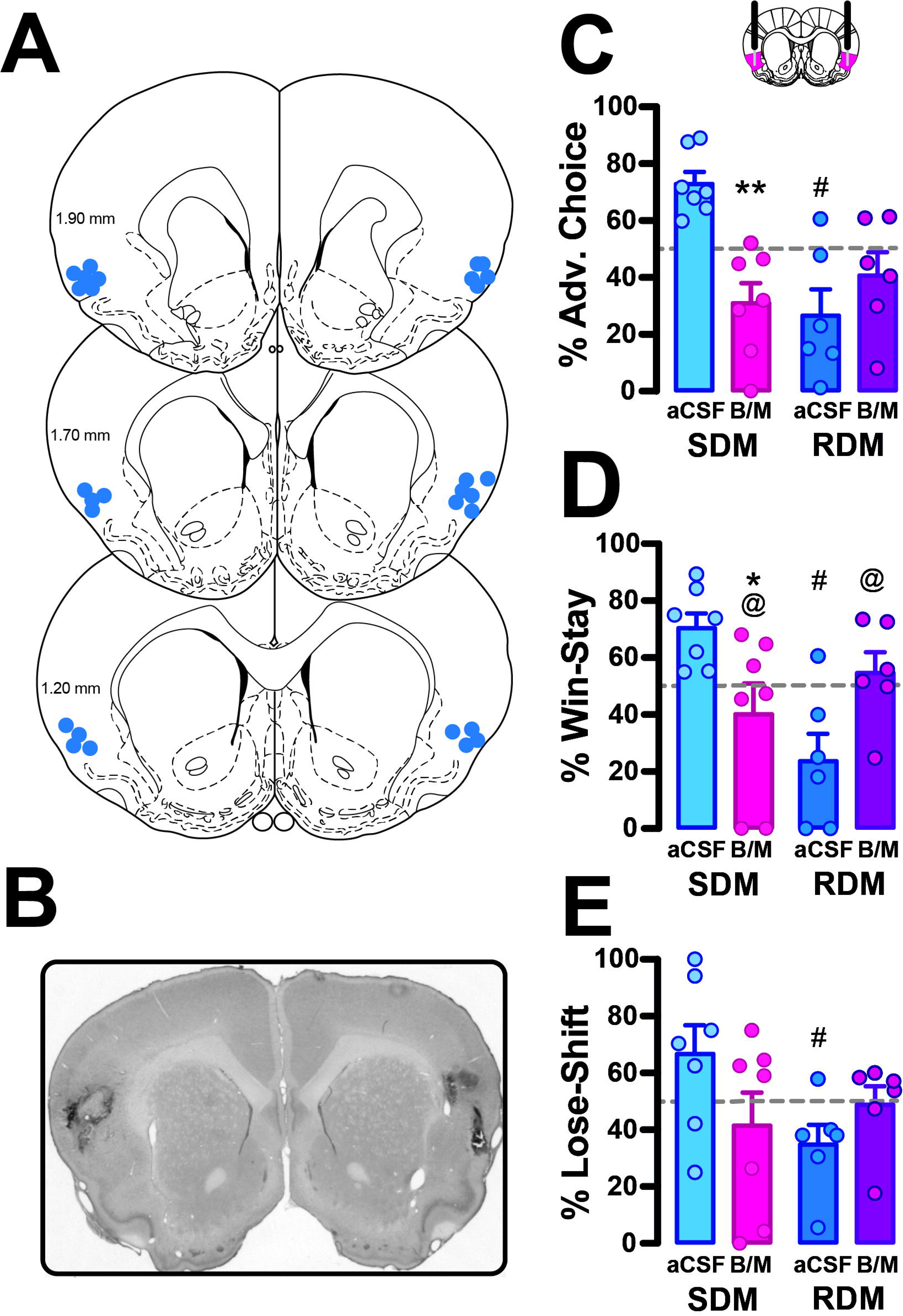
Inactivation of the AIC improves decision-making performance of RDMs and worsens that of SDMs by altering responsivity to reward. (**A**) Histologically confirmed anatomical locations of the injector tip for all bilateral AIC inactivations. (**B**) Representative 5X micrograph of a cresyl violet- stained coronal brain section used to confirm the location of the tip of the injector targeting the AIC. (**C**) AIC inactivation reduced the frequency of advantageous choices in SDMs. (**D**) AIC inactivation decreased the win-stay probability in SDMs while increasing it in RDMs. (**E**) AIC inactivation had no effect on loose-stay behaviour. * and ** indicate p < 0.05 or p < 0.01 for aCSF vs. BM within a phenotype, respectively. #: aCSF-infused RDMs differed from aCSF-infused SDMs. @ denotes where inactivation causes performance to reach chance level.

**Figure 3.**
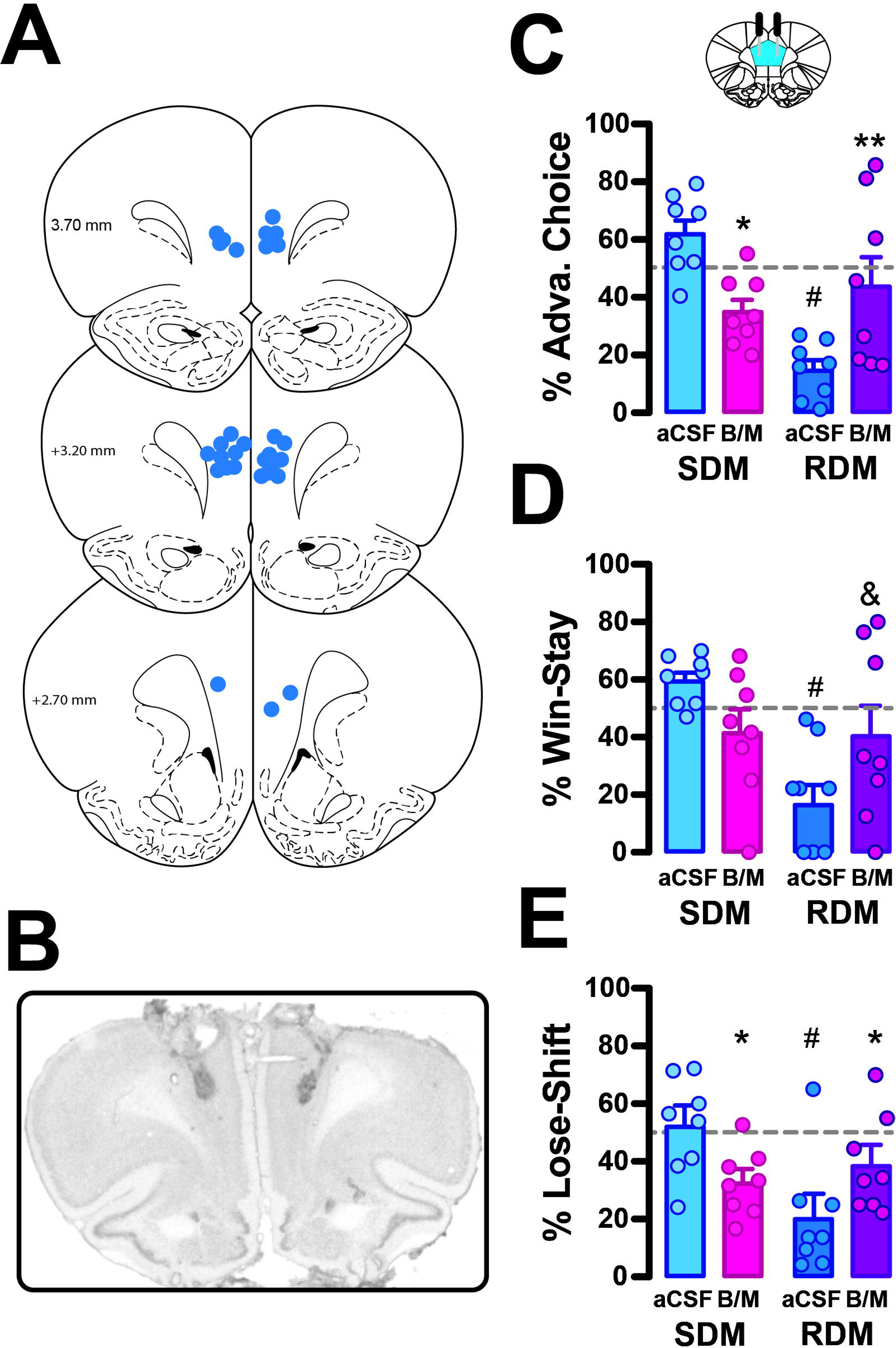
PRL inactivation drives alterations in loss responsivity that improve decision-making performance of RDMs and worsen that of SDMs. (**A**) Histologically confirmed anatomical locations of the injector tip for all bilateral PLC inactivations. (**B**) Representative 5X micrograph of a cresyl violet- stained coronal brain section used to confirm the location of the tip of the injector targeting the PRL. (**C**) PRL inactivation resulted in an increase in the frequency of advantageous choices in RDMs and a decrease in SDMs. (**D**) PRL inactivation had no effect on the win-stay behaviour of SDMs and tended to increase it in RDMs. (**E**) PRL inactivation decreased the win-stay probability in SDMs while increasing it in RDMs. * and ** indicate p < 0.05 or p < 0.01 for aCSF vs. BM within a phenotype, respectively. & indicates trend-level effect of inactivation within a phenotype (p = 0.054). # indicates significant differences between phenotypes at aCSF baseline.

### Local inactivation

Ten days after the surgery, rats used in experiments 1 and 2 underwent two other rGT test sessions following bilateral aCSF or B/M infusions (counterbalanced across animals), in which contingencies were systematically reshuffled to prevent carryover effects from the previous testing session [exactly as in 5].

For experiment 3, three test sessions were conducted, with contralateral or control ipsilateral inactivations of the AIC and PRL, in a counterbalanced design, which were both compared to the same aCSF test condition. For each of these test sessions, contingencies were shuffled differently from one another and from the original phenotyping session.

Before these test sessions, a 28-gauge stainless steel hypodermic injector (Plastics One, Roanoke, VA, USA) was lowered into each guide cannula to protrude 2 mm from its ventral end, and a cocktail of baclofen and muscimol (0.03 nmol/0.5 µL/infusion) or the same volume of Vehicle (phosphate-buffered saline, pH 7.4) was infused over 90s using a syringe pump (Harvard Apparatus, Holliston, MA, USA), followed by a 60-s diffusion period. The injector was then removed, the obturator replaced, and behavioural test sessions began 5 minutes later.

For all experiments, between test sessions, rats were trained in several forced-choice sessions to avoid carryover effects. Before these RGT challenges, they were habituated to the insertion of the injectors and infusions. First, an injector was lowered into each guide cannula to protrude 2 mm from its ventral end and then, two days later, on two separate occasions, three days apart, each rat received two infusions of vehicle into each side of the brain.

### Histology

At the end of the experiment, rats were euthanised with an overdose of sodium pentobarbital (300 mg; Dolethal; Vétoquinol UK Ltd, Buckingham, UK), and perfused transcardially with isotonic saline, followed by 10% neutral buffered formalin. Brains were extracted and transferred to a 30% sucrose solution in 0.01 M PBS for 48 hrs before sectioning at 60 μm using Leica CM3050 S cryostat. Sections were mounted and stained with Cresyl Violet as previously described [35]. Cannulae placements in the PLC and AIC were verified using a light microscope, which led to the post-hoc exclusion of 3 rats (AIC: 1 SDM and 2 RDMs) (**Figures 2A, 2B**, 3A, 3B**, 4A, and 4B**).

**Figure 4.**
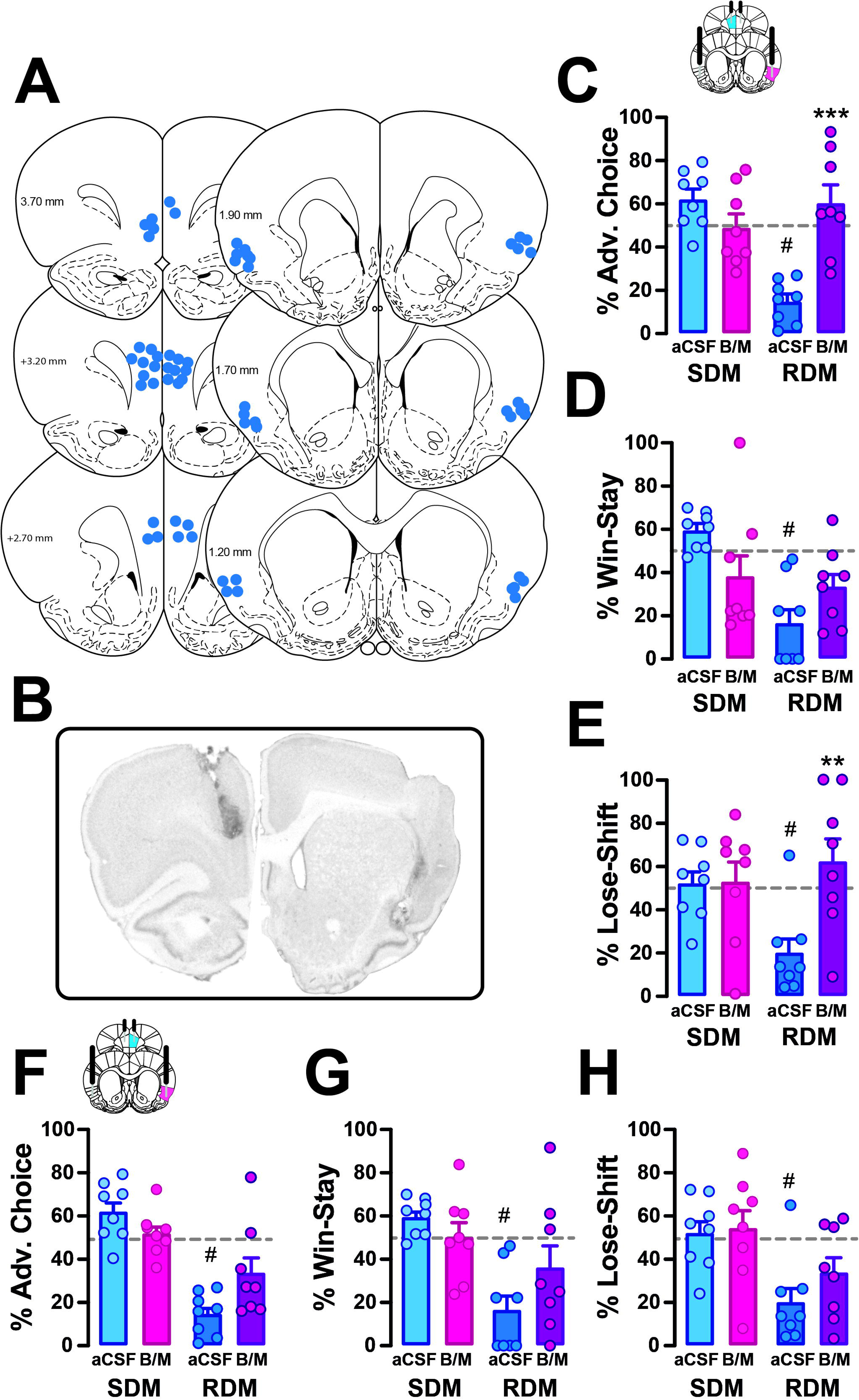
Functional disconnection of the PRL and AIC selectively improves decision making in RDMs by enhancing loss sensitivity. (**A**) Histologically confirmed anatomical locations of the injector tip for all bilateral AIC + PRL inactivations. (**B**) Representative 5X micrograph of a cresyl violet- stained coronal brain section used to confirm the location of the tip of the injectors targeting the PRL and AIC. (**C**) Functionally disconnecting the PRL and the AIC increased the frequency of advantageous choices in RDMs but had no effect in SDMs. (**D**) Functionally disconnecting the PRL and the AIC had no effect on win-stay behaviour. (**E**) Only RDMs exhibited an increase in lose-shift behaviour following functional disconnection of the PRL and AIC. (**F**) Simultaneous ipsilateral inactivation of the PRL and the AIC had no effect on the frequency of advantageous choice, (G) win-stay behaviour, or (H) lose-shift behaviour displayed by SDMs and RDMs. ** and *** indicate p < 0.01 or p < 0.001 for aCSF vs. BM within a phenotype, respectively. # indicates significant differences between phenotypes at aCSF baseline.

### Data and Statistical analyses

Data were analysed and plotted using Prism for macOS (GraphPad Software, USA). Assumptions of normality, homogeneity of variance, and sphericity were assessed using the Shapiro–Wilk, Levene, and Mauchly sphericity tests, respectively. Repeated-measures analysis of variance (RM-ANOVA) with treatment (VEH vs. BM) as the within-subjects factor and phenotype (SDM vs. RDM) as the between-subjects factor was used to detect omnibus effects. Independent samples t-tests were used to verify that SDMs differed from BDMs on all behavioural measures. Significant RM-ANOVAs were followed up with planned Fisher’s LSD comparisons of VEH vs BM for each phenotype or one-sample t-tests to determine whether decision-making strategy differed from chance. For all analyses, the α criterion was set at p = 0.05. Effect sizes are reported as partial eta squared (pη2).

## Results

### Experiment 1 – Inhibition of the anterior insular cortex (AIC)

Bilateral AIC inhibition resulted in a bidirectional effect on decision-making performance in SDMs and RDMs [phenotype x inactivation interaction: F_1,11_ = 18.69, p = 0.001, pη2 = 0.310] (**Figure 2C**), predominantly mediated by an effect on win-stay tendencies [phenotype x inactivation interaction: F_1,11_ = 10.58, p = 0.008, pη2 = 0.332] (**Figure 2D**). In SDMs, the baseline performance of which was much better than that of RDMs [RDM_aCSF_ vs. SDM_aCSF_: t_11_ = 4.79, p < 0.001] (**Figure 2A**), bilateral AIC inhibition reduced the tendency to choose advantageous options by decreasing win-stay tendencies, suggesting a worsening of decision-making in the gain domain. The same manipulation slightly increased performance in RDMs (**Figure 2C**) by increasing their win-stay tendency (**Figure 2D**). Consequently, while baseline win-shift behaviour in RDMs was significantly lower than in SDMs [RDM_aCSF_ vs. SDM_aCSF_: t_11_ = 4.27, p = 0.003], AIC inhibition abolished this difference [t_11_ = −1.11, p = 0.294], bringing win-shift behaviour to chance level in both groups [SDM _win-stay ≠ 50_: t_7_ = −0.88, p = 0.411; RDM _win-stay ≠ 50_: t_6_ = 0.67, p = 0.534]. In contrast, the tendency to sample a different option after a negative outcome, in which SDMs also differed from RDMs at baseline [RDM_aCSF_ vs. SDM_aCSF_: t_11_ = 2.51, p = 0.026], was not affected by AIC inactivation [main effect of phenotype: F_1,11_ = 2.968, p = 0.113, pη2 = 0.059; inactivation: F_1,11_ = 0.248 p = 0.628, pη2 = 0.012; phenotype x inactivation interaction: F_1,11_ = 3.01, p = 0.111, pη2 = 0.15] (**Figure 2E**).

### Experiment 2 – Inhibition of prelimbic cortex (PLC)

As with the AIC, bilateral PLC inhibition had a bidirectional effect on decision-making performance in SDMs and RDMs, which also differed from one another at baseline [phenotype x inactivation interaction: F_1,14_ = 18.99, p < 0.001, pη2 = 0.350; RDM_aCSF_ vs. SDM_aCSF_: t_14_ = 8.22, p < 0.001], decreasing the probability of choosing the advantageous option in SDMs and increasing it in RDMs (**Figure 3C**). However, in contrast with those of AIC inhibition, these effects were driven largely by changes in lose-shift behaviour [phenotype x inactivation interaction: F_1,14_ = 11.50, p = 0.004, pη2 = 0.244], on which the phenotypes differed at baseline [RDM_aCSF_ vs. SDM_aCSF_: t_14_ = 3.50, p = 0.004 (**Figure 3E**). In win-stay behaviour, which also differed between phenotypes at baseline [RDM_aCSF_ vs. SDM_aCSF_: t_14_ = 5.65, p < 0.001], PRL inactivation caused a marginal increase in RDMs only [phenotype x inactivation interaction: F_1,14_ = 6.820, p = 0.021, pη2 = 0.173; RDM_aCSF_ vs. RDM_BM_: p = 0.054] (**Figure 3D**).

### Experiment 3 – Functional disconnection of the AIC and PLC

At baseline, SDMs and RDMs differed on all behavioural measures [all ps < 0.02]. Functional disconnection of the PLC and the AIC exclusively influenced the performance of RDMs [phenotype x disconnection interaction: F_1,14_ = 23.760, p < 0.001, pη = 0.350] in whom the manipulation resulted in a substantial improvement in performance while having little effect on SDMs’ (**Figure 4C**). This improvement in the decision-making ability of RDMs following a functional disconnection of the PLC and the AIC was not due to any effect on win-stay tendencies [phenotype x disconnection interaction: F_1,14_ = 0.077, p = 0.785, pη2 = 0.021] (**Figure 4D**), but to an increase in lose-shift behaviour [phenotype x Treatment: F_1,14_ = 5.158, p < 0.039, pη = 0.136) (**Figure 4E**). Following functional disconnection of the PLC and the AIC, performance in each of these measures was greatly increased in RDMs so much so that it no longer differed from that of SDMs at baseline (**Figure 4C-E**).

These effects were specific to the functional disconnection between the PLC and the AIC, as double ipsilateral inactivation of the PLC and AIC did not substantially alter performance in either group, the small interactions found being primarily driven by the low level of performance of the aCSF-infused SDM group [phenotype × inactivation interaction for both overall choice of advantageous options: F_1,14_ = 6.714, p = 0.0213, pη2 = 0.104, win–stay behaviour: F_1,14_ = 4.777, p = 0.0463, pη2 = 0.081) and lose–shift behaviour: F_1,14_ = 0.5124, p = 0.4859, pη2 = 0.014] (**Figure 4F-H**).

## Discussion

The present findings demonstrate that the AIC and PLC independently exert opposite effects on probabilistic decision-making in SDMs and RDMs, supporting good performance in the former and hindering it in the latter, with the AIC and PLC controlling predominantly win-stays and lose shifts, respectively. Most importantly, while functionally disconnecting the AIC and the PLC has no effect on decision-making in SDMs, it substantially increased performance in RDMs. Thereby, this demonstrates that individual differences in probabilistic decision making are underlined by distinct functional interactions between the AIC and the PLC, which are connected by reciprocal, mostly unilateral, projections [36–38]

The results of experiment 1 replicated and extended previous findings demonstrating that irreversible bilateral excitotoxic lesions to the AIC produce phenotype-dependent effects on decision-making, improving performance in RDMs while impairing it in SDMs [5], a deficit predominantly due to an inability to switch from exploration to exploitation. Here, trial-by-trial analyses revealed that this effect was predominantly driven by alterations in win–stay behaviour, which was profoundly reduced in SDMs and greatly increased in RDMs after bilateral AIC inactivation. This observation suggests that the AIC may differentially encode an interoceptive representation of the incentive value of positive outcomes (i.e., rewards) in SDMs and RDMs.

In SDMs, this function appears to be adaptive: the AIC amplifies the impact of positive outcomes, promoting stable exploitation of advantageous options, in line with evidence that, in humans, the insula exhibits heightened activity during a decision following a win compared to a loss [39]. Transient inactivation of the AIC results in the loss of this amplification, resulting in a weakening win–stay behaviour and a degradation of performance. In contrast, in RDMs, the AIC may convey an interoceptive signal of poor fidelity in response to a win that is noisy and/or ambiguous, preventing learning from interoceptive representations of the outcome, the removal of which, following post-training lesion or inactivation of the AIC, results in a less noisy system and improved performance.

This theoretical framework is further supported by evidence from experiment 1 that, following AIC inactivation, win–stay behaviour regressed to chance levels in both SDMs and RDMs, indicating that the AIC exerts symmetrically opposite modulatory roles on the salient or reinforcing properties of positive outcomes under uncertainty in these groups.

At the neurochemical level, AIC-mediated coding of the incentive value of positive outcomes during decision making under uncertainty has been shown to be dependent on Dopamine D2 receptors [40], a mechanism that may underlie dopamine control of decision making performance in the RGT as revealed by chemogenetic manipulation of VTA dopamine neurons [41,42] or systemic dopamine receptor agonism [43].

Experiment 2 revealed a complementary yet distinct role for the PLC in decision-making under uncertainty. In contrast to AIC inactivation, which selectively affected win–stay behaviour, PLC inactivation predominantly altered responses to losses. Specifically, PLC inactivation increased lose–shift behaviour in RDMs, improving their performance, while reducing loss sensitivity in SDMs and hence impairing their performance. These findings indicate that the PLC contributes to behavioural adaptation following losses, likely by processing negative feedback to adjust decision-making policy, in line with previous demonstrations of the role of the PLC in the evaluation of negative outcomes, error monitoring, and the subsequent implementation of behavioural control [44–49]. Lesions to, or inactivation of this region have been associated with impaired decision-making [29,50], the functional equivalent of which has been shown to be less activated in patients with gambling disorders or substance use disorder during the manifestation of their decision making impairment in IGT [18,51].

Within this framework, the PLC may function as a negative-outcome detector, transforming losses into behavioural-change signals. Critically, the present data suggest that this process is not uniform across individuals. In SDMs, PLC activity supports adaptive flexibility, promoting switching following losses. When the PLC is inactivated, this capacity is diminished, leading to perseverative behaviour even in the face of negative outcomes. In RDMs, however, PLC function appears to be constrained or distorted, limiting sensitivity to losses. Inactivation of the PLC in these individuals paradoxically enhances loss-driven switching, suggesting that the intact PLC may contribute to maladaptive post-loss persistence in this group.

A central question arising from these findings is how the AIC and PLC interact to shape behaviour in SDMs and RDMs. The functional disconnection carried out in experiment 3 provides critical insight in this regard. Functionally disconnecting the AIC and the PLC selectively improved performance in RDMs by increasing lose–shift behaviour, while leaving SDMs unaffected. This AIC-PLC disconnection had a more profound effect on lose-shift behaviour than the PLC inactivation alone, suggesting that the tendency to engage in risky decision-making exhibited by RDMs depends on coordinated (albeit maladaptive) activity between the two regions. This observation resonates with functional neuroimaging evidence of enhanced co-recruitment of the insula and prefrontal regions (including the vmPFC) during decision-making in individuals with a gambling disorder [52].

These findings support a model in which the AIC and PLC can operate either in parallel or in series across individuals, thereby shaping individual tendencies to adopt safe vs risky decision-making, respectively. In SDMs, decision-making appears to rely on parallel processing: the AIC encodes the incentive value of rewards, promoting behavioural stability, while the PLC independently processes negative outcomes, enabling adaptive switching. Because these systems operate relatively independently, disrupting their interaction has minimal impact on behaviour. In contrast, in RDMs AIC–PLC functional connectivity appears to behave in a more serial and interdependent manner. In this configuration, noisy AIC-generated interoceptive signals could be transmitted to the loss-tuned PLC, where interoceptive signals of gain could be attributed with negatively-valenced incentive value. Conversely, PLC-generated evaluations of loss may be communicated to a gain-biased AIC, misattributing the events with positive incentive valence. Disrupting this circuit effectively frees the PLC from insula-driven noise, allowing it to more accurately evaluate negative outcomes and promote adaptive behavioural change. This framework provides a mechanistic account of why disconnection selectively benefits RDMs. It is not simply that either region is dysfunctional in isolation, but rather that their interaction reinforces maladaptive processing. In SDMs, where signals are well-calibrated, such coupling is either absent or functionally benign for updating decision-making policies (**Figure 5**).

**Figure 5.**
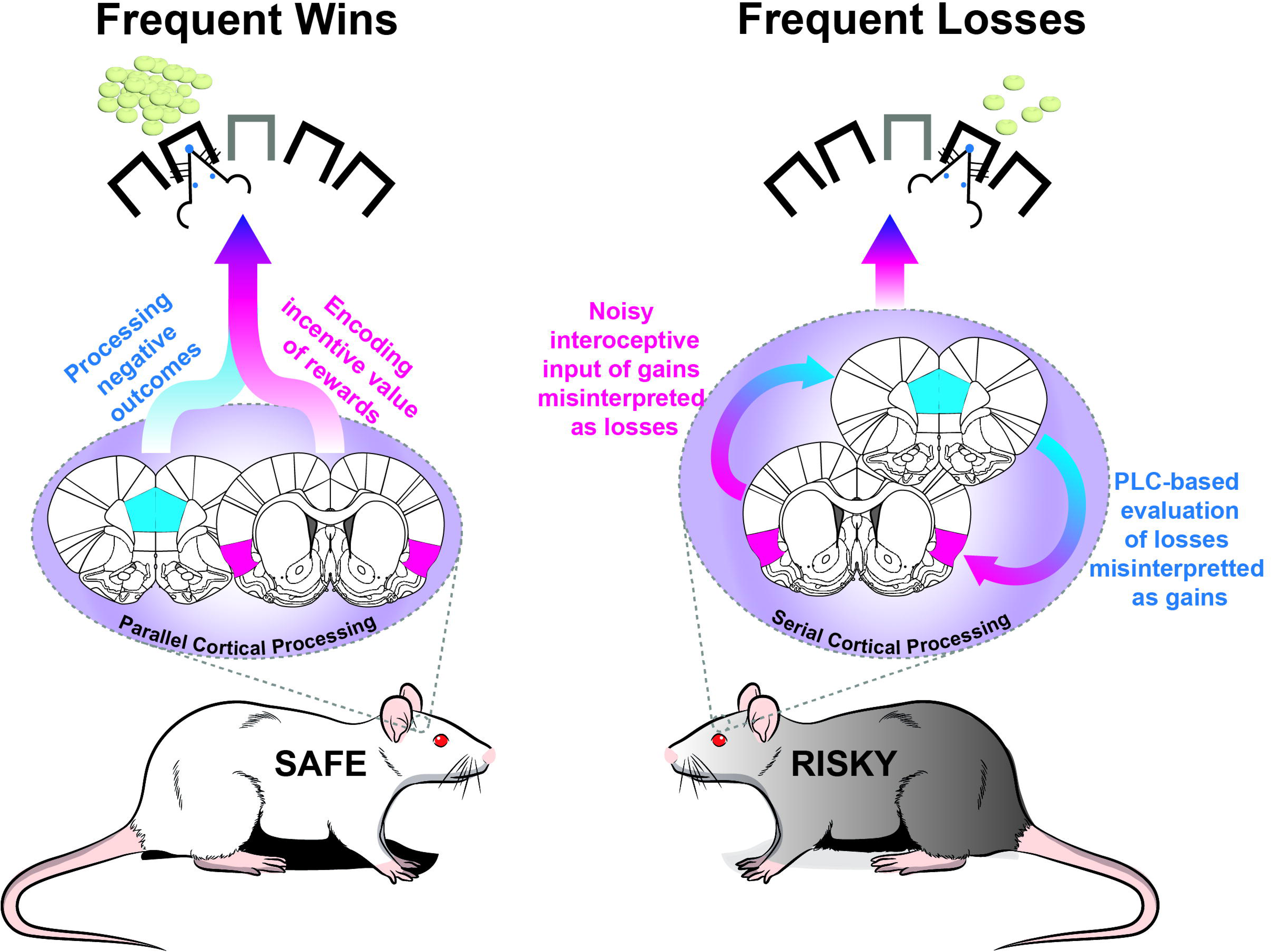
Parallel vs serial processing of wins and losses by the AIC-PLC network underlies SDM vs RDM phenotype-related differences in decision making. In this graphical summary, we speculate that in SDM rats, the AIC encodes the incentive value of rewards, in parallel with the processing of the valence of losses in the PLC. In contrast, in RDM rats, the AIC and PLC process information serially. Risky decision-making can be speculated to be predicated either on aberrant integration by the AIC of PLC-encoded losses as gains, or on noisy interoceptive signals integrated by the AIC from gains, which are misinterpreted as losses upon subsequent integration in the PLC.

The present findings may have important implications for understanding the neurobiological mechanisms underlying gambling disorder (GD), particularly how poor interoceptive sensitivity contributes to maladaptive reward and loss processing in GD [53–55]. Under normal conditions, losses during gambling should evoke aversive interoceptive states, such as tension, discomfort, frustration, or autonomic arousal, which serve as negative teaching signals that promote behavioural adjustment and discourage maladaptive persistence [24,39]. However, individuals with GD may exhibit reduced sensitivity to these interoceptive signals, impairing their ability to appropriately register the negative emotional and bodily consequences of losing [54,56].

Such interoceptive deficits may be particularly important in the context of modern gambling environments that exploit losses disguised as wins and near misses. Although losses disguised as wins are objectively negative outcomes, they are often paired with salient audiovisual feedback that resembles genuine reward, producing physiological and subjective responses more similar to wins than losses [57,58]. Likewise, near misses can generate heightened arousal and motivate continued gambling despite the absence of reward, particularly in vulnerable individuals who interpret these outcomes as signals of being “close” to winning [59,60], a cognitive distortion that depends on the insula [59]. If individuals with GD are less sensitive to the aversive interoceptive consequences of losing, these misleading outcomes may fail to generate the negative affective states necessary for adaptive behavioural updating. Consequently, gambling behaviour may become increasingly guided by inaccurate interoceptive representations of positive and negative outcomes. The present findings suggest that the AIC may contribute to this process by modulating the interoceptive representation of gambling outcomes, while the PLC integrates this information to guide and update internal rules under uncertainty in response to losses.

In risky decision makers, a serial functional coupling between the PLC and the AIC, instead of a parallel engagement of these regions may facilitate distorted interoceptive signals to bias prefrontal evaluation of outcomes, thereby weakening loss sensitivity and promoting maladaptive persistence despite repeated adverse consequences.

In conclusion, the results of the present study reveal that the AIC and the PLC independently contribute to decision-making, thereby maximising gain under uncertainty in individuals with a tendency to make safe decisions through parallel processing, whereas their functional interaction underlies suboptimal choices in individuals with a tendency to make risky decisions.

## Acknowledgements

This work was carried out at the Departments of Psychology and Physiology, Development and Neuroscience of the University of Cambridge and was supported by UKRI (Medical Research Council) grants to DB [MR/W019647/1] and Barry Everitt, DB et al. [MR/N02530X/1]. DJ was supported by a BBSRC-DTP/Shionogi joint grant to DB (G101457). KJ was supported by an Early Post Doc mobility Fellowship (2021-2022) from the Swiss National Science Foundation. TH was supported by Leverhulme & Isaac Newton Trust early career fellowships, and a new faculty start-up grant from Simon Fraser University.

## Author contributions

DB and KJ designed the experiment. DJ wrote the custom Python scripts to extract and compile the raw behavioural data. KJ and DJ performed the behavioural experiments. KJ and LS conducted the histology. TH and KJ analysed the data. DJ TH and DB wrote the manuscript.

